# Reentrant condensation of a multicomponent complex system of biomolecules induced by polyphosphate

**DOI:** 10.1101/2023.03.02.530750

**Authors:** Tomohiro Furuki, Tomohiro Nobeyama, Shunji Suetaka, Ryokei Matsui, Tatsuhiko Fukuoka, Munehito Arai, Kentaro Shiraki

**Author notes:** Corresponding author. E-mail address (K.Shiraki).

## Abstract

Reentrant condensation (RC) is a phase behavior of protein solution comprising at least two components. In RC, a protein state varies from one phase to two phases and then back to one phase as the concentration of one component monotonically increases. To understand the phase behavior of multicomponent complex solutions of biomolecules, it is worth constructing an experimental multicomponent system that exhibits RC behavior. Here, we used a cola/milk mixture to investigate RC of a multicomponent complex system and explained the RC mechanism by reducing the system to two pure components, polyphosphate (polyP) and casein. In the multicomponent complex system, RC was observed with 20–60% cola and 1% milk. In the pure system, RC occurred with 0.01–2 mM tetraphosphate and 0.5 mg/ml casein. Moreover, the phase diagram showed that the condensation of casein depended on the chain length of the polyP. The present study succeeded in experimentally inducing RC in a multicomponent system and reproducing RC even when the system was reduced to its pure components. The fact that RC can be experimentally induced using common materials will provide important insights into the understanding of phase-separation behavior of biomolecules.

## 1. Introduction

Proteins have diverse states in solution. Developing methods for controlling a protein state in solution would benefit various fields, including the food industry and pharmaceutical sciences [1]. Cytosols are highly concentrated protein solutions, and the protein phase behavior, such as liquid–liquid phase separation (LLPS), is implicated in various biological phenomena [2, 3]. Proteins in solution are considered to have two states: a one-phase state in which protein molecules are homogeneously dispersed and a two-phase state in which protein molecules are heterogeneously dispersed (typically with condensation, aggregation, or phase separation). Coagulants are additives that change the protein solution from the one-phase state to the two-phase state (Fig. 1a). For example, CaSO_4_ turns soy milk into tofu (soybean curd) [4]. Similarly, the addition of polyglutamic acid to an antibody solution induces the solution to separate into two phases by forming a stable protein–polyelectrolyte complex (PPC) [5]. In contrast, aggregation suppressors are additives that prevent a protein solution from separating into two phases, thus maintaining proteins dispersed in one phase (Fig. 1b). For example, 0.5 M NaSCN completely prevents thermal aggregation of hen egg-white proteins and keeps them in one phase even at 90ºC [6]. Additionally, some additives can both promote and inhibit condensation, depending on their concentration. For example, arginine can prevent antibody aggregation and can also promote the formation of soluble aggregates [7].

**Fig. 1.**
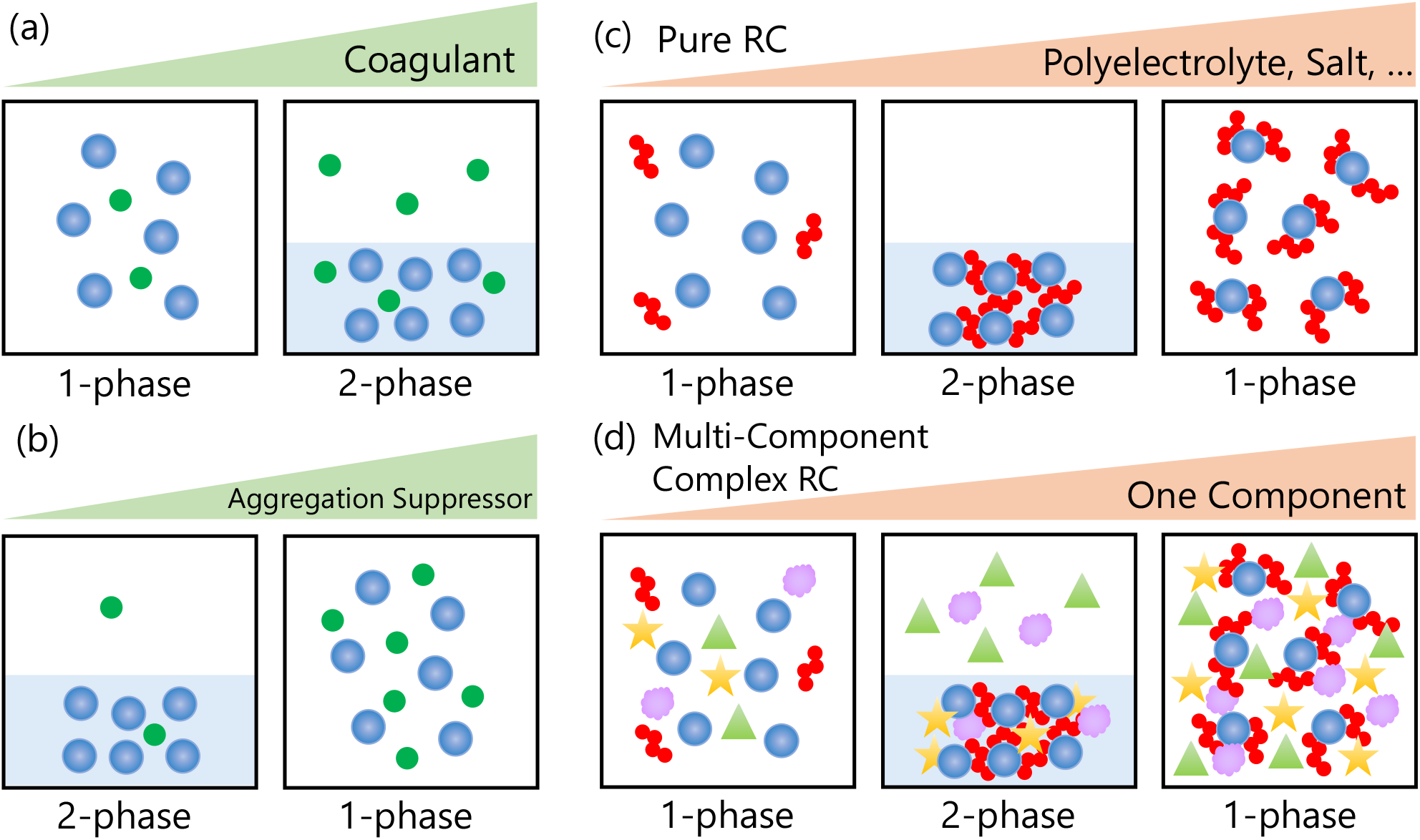
Schematic models of various phase behaviors of protein solutions. (a) Coagulants (green) induce condensation of proteins (blue). (b) Aggregation suppressors (green) inhibit the condensation of proteins (blue). (c) RC in a two-component pure system consisting of a protein (blue) and a polyelectrolyte or salt (red). (d) RC in a multicomponent complex system consisting of a protein (blue), a polyelectrolyte or salt (red), and other components (other colors). In (c) and (d), an increase in the concentration of one component (e.g., polyelectrolyte) induces RC.

Reentrant condensation (RC) is a phenomenon in which a system changes from a one-phase state to a two-phase state and then returns to a one-phase state as the concentration of one component in the system monotonically increases (Fig. 1c). The one-phase states at low and high concentrations of the component are macroscopically identical but differ at the molecular level. Recently, RC has been widely observed in colloidal systems [8–11] and in proteins and polypeptides [12–17]. When Na^+^, Mg^2+^, Al^3+^, and Zr^4+^ are added to silica nanoparticle solutions, trivalent Al^3+^ and tetravalent Zr^4+^ can induce RC, whereas monovalent Na^+^ and divalent Mg^2+^ cannot [8]. Trivalent cations, such as Y^3+^, La^3+^, Fe^3+^, and Al^3+^, induce RC of globular proteins such as bovine serum albumin (BSA), human serum albumin, and ovalbumin because ions with high charge density can easily induce attractive interactions between ions and proteins [12, 13]. The mechanism of RC in protein solutions is as follows. Proteins are negatively charged at pH values above their isoelectric points (p*I*). Oppositely charged cations cancel out the charges of proteins, reducing the repulsion between proteins and polyelectrolytes, thereby inducing condensation. Further addition of cations reverses the charges on the protein surface, resulting in repulsive forces and the system returning to one phase [12, 13]. The occurrence of such charge inversion is supported by molecular dynamics simulations in a tetra-aspartate/trivalent cation system [14]. RC is also observed with other biomolecule combinations, such as RNA/positively charged peptides [15] and intrinsically disordered proteins/salts [16, 17]. The common feature of these two-component systems exhibiting RC behavior is that one or both components are multivalent. Recent studies have suggested that RC contributes to biochemical timekeeping by generating condensates that form and dissolve as the concentration of transcription products monotonically increases [18]. However, RC has not been directly observed in living cells because it is impossible to monotonically change the concentration of only one component in living cells. It is also unclear whether RC occurs in a multicomponent complex system and, in which case, whether it can be reproduced using the pure components of the system (Fig. 1d). Therefore, generating an artificial model system that exhibits RC behavior and consists of multiple complex components, and then reconstructing the system using its pure components would allow to apply the knowledge obtained so far in pure systems.

Polyphosphate (polyP) is a negatively charged linear polymer consisting of three to several thousand inorganic phosphate units. PolyP is highly conserved in living organisms since ancient times and has been identified in every organism tested [19, 20]. At acidic pH, polyP interacts with positively charged proteins through electrostatic interactions, thus promoting aggregation and amyloid formation [21, 22]. At neutral pH, polyP behaves as kosmotrope and dehydrates water molecules around β_2_-microglobulin to accelerate β_2_-microglobulin aggregation and amyloid formation [21]. Depending on their chain length, PolyP also induces LLPS of positively charged green fluorescent protein (+36GFP) [23]. Condensation of polyP, DNA, and Hfq protein leads to heterochromatin formation in bacteria [24]. In addition, polyP is widely used as an industrial reagent in water treatment, fertilizers, and flame retardants [25]. PolyP is also used as a food additive because of its water retention, antibacterial, and buffering properties [25]. In cola, polyP refines carbon dioxide bubbles and maintains them even after 27 days of storage [26].

Interestingly, polyP induces RC behavior of biomolecules. In cheese production, polyP promotes casein aggregation at neutral pH [27], whereas an excess of polyP suppresses casein aggregation at neutral pH by imparting a large repulsive negative charge on the casein [28]. Furthermore, polyP induces RC of histatin 5 and lysozyme due to their high density of negative charges [29, 30]. In this case, arginine interacts more strongly with polyP than lysine, although they have the same positive charge [29]. At moderate concentrations, triphosphate (triP) forms gels with positively charged polyarylamines through electrostatic interactions, whereas higher concentration of triP dissolves the gels by reversing the surface charge of the triP/polyarylamine complex [31]. Thus, polyP can be used as a component of a multicomponent complex system that exhibits RC behavior.

In the present study, we used a cola/milk system to investigate whether RC can occur in a multicomponent complex system and whether the mechanism of RC can be explained on the basis of its pure components. Cola and milk are models of multicomponent complex systems containing polyP and casein, respectively. It is well known that mixing cola and milk induces condensation [32]. Strikingly, we observed RC with 20–60% cola and 1% milk. As a pure system, we used casein and polyP of different chain lengths and observed RC with 0.01–2 mM tetraphosphate (tretaP) and 0.5 mg/ml casein. This is the first experimental demonstration of RC in a multicomponent complex system, the mechanism of which could be explained by reducing the system to its pure components. This study provides fundamental knowledge for understanding the phase behavior of multicomponent complex systems.

## 2. Materials and methods

### 2.1. Materials

Sodium caseinate (casein) was purchased from Wako Pure Chemical Industries (Tokyo, Japan). Sodium tetraP, triP, and diphosphate (diP) were purchased from Nacalai Tesque (Kyoto, Japan). Pepsi cola was purchased from Suntory (Osaka, Japan). Milk was purchased from Meiji (Tokyo, Japan). Casein and each polyP were dissolved separately in Milli-Q water. Cola was used after removing carbon dioxide bubbles by sonication. Sample solutions were made by mixing the solutions at the desired concentrations. The pH of each sample was adjusted by adding HCl or phosphoric acid and NaOH (up to 10 mM).

### 2.2. Turbidity measurement

Turbidity was analyzed to detect the color of cola and turbidity of milk. Sample solutions were made and centrifuged (18,800×*g*, 20 min, 20 ºC). Then, the supernatant turbidity was determined by measuring the absorption at 400 nm using a NanoDrop ND-1000 (NanoDrop Technologies, Delaware, US) at room temperature.

### 2.3. Casein precipitation analysis

Casein precipitation was analyzed to determine casein concentration in the supernatant of the sample solutions. Sample solutions were made and centrifuged (18,800×*g*, 20 min, 20 ºC). Then, casein concentration in the supernatant was determined using the Bradford or Lowry methods. The Bradford protein assay kit was purchased from Takara Bio (Shiga, Japan). Aliquots (4 µl) of supernatant samples were placed in 96-well clear flat bottom ultra-low attachment microplates (Corning, New York, US), and 200 µl of Bradford dye reagent were added. After incubation for 30 min, the absorption at 595 nm was measured for each sample using a microplate reader (Infinite 200 PRO, Tecan Japan, Kanagawa, Japan) at 25 ºC. The measurements were performed in triplicate. The determination of casein concentration by the Lowry method was performed using the DC™ protein assay kit purchased from Bio-Rad Laboratories (California, US). Aliquots (5 µl) of supernatant samples were placed in 96-well microplates, and 25 µl of the DC protein assay reagent A and 200 µl of reagent B were added. After incubation for 30 min, the absorption at 750 nm was measured for each sample using a microplate reader at 25 ºC. The measurements were performed in triplicate. For each method, a calibration curve was generated using pure casein solutions and was used to determine casein concentration.

### 2.4. Zeta potential measurement

Zeta potential was measured on a Zetasizer Nano ZS using DTS1070 cells (Malvern Instruments, Malvern, UK) at 25 ºC. Samples containing 0.5 mg/ml casein and 0.01–2 mM tetraP were centrifuged (18,800×*g*, 20 min, 20 ºC), and the supernatants were collected the day before zeta potential measurements.

### 2.5. Generation of phase diagrams of casein in polyP solutions

Phase diagrams of casein in polyP solutions were generated using precipitation measurements. The solutions were mixed and centrifuged (15,000 rpm, 30 min, 4 ºC). Then, the absorption of the sample supernatants at 280 nm was measured on a UV-1800 spectrophotometer (Shimadzu, Kyoto, Japan) at room temperature. A calibration curve obtained using pure casein solutions was used to determine casein concentration. When more than 90% of casein was precipitated, we considered it to be “condensed,” whereas casein was considered “soluble” when less than 90% was precipitated.

## 3. Results

To understand the phase behavior of a cola/milk mixture (Fig. 2a), we investigated the cola concentration dependence of the supernatant turbidity. Solutions at pH 3.2–3.6 and 7.0–7.4 were prepared with a constant concentration of milk (1%) and different concentrations of cola (0–97%). Fig. 2b shows the supernatant turbidity of these solutions. At pH 3.2–3.6, the supernatant turbidity decreased to around 0 in the presence of 30–40% cola, whereas it was 0.25–0.50 for other cola concentrations. In contrast, at pH 7.0–7.4, the supernatant turbidity was 0.26–0.38 for all cola concentrations tested. Fig. 2c shows the ratio of the supernatant turbidity at pH 3.2–3.6 to that at pH 7.0–7.4. In the presence of 30–40% cola, the ratio was below 0.07, whereas it was above 0.95 with other cola concentrations. Interestingly, at pH 3.2–3.6, the supernatant turbidity first increased, then decreased, and finally increased again as the cola concentration increased. These data indicated that RC occurred in the cola/milk system.

**Fig. 2.**
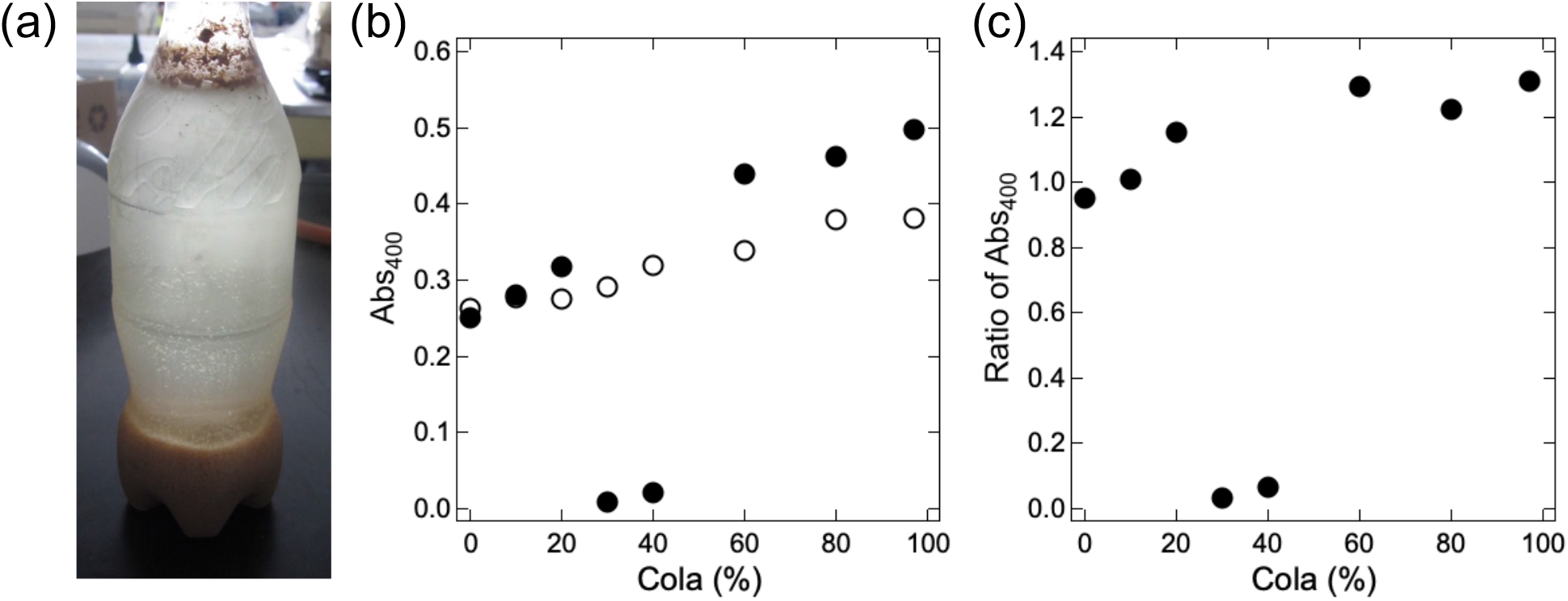
RC in the cola/milk system. (a) Representative picture of the cloudy white milk precipitates within dark brown-colored cola, resulting in a clear supernatant. Movies of the cola/milk mixture can be found on YouTube [32]. (b) Supernatant turbidity of a mixture of 1% milk and different concentrations of cola at pH 3.2–3.6 (closed circles) and pH 7.0–7.4 (open circles). (c) Ratio of the supernatant turbidity at pH 3.2–3.6 to that at pH 7.0–7.4 in the cola/milk system.

Next, casein was selected among the milk components because it is the major milk protein. To understand the phase behavior of the cola/casein mixture, we investigated the effects of the cola concentration on casein concentration in the supernatant and the supernatant turbidity. Solutions were prepared using a constant concentration of casein (0.5 mg/ml) and varying concentrations of cola (0–97%) at pH 3.2–3.6 and 7.0–7.4. Fig. 3a shows casein concentration in the supernatant of cola/casein mixtures containing different concentrations of cola. As the cola concentration increased from 0% to 60%, casein concentration in the supernatant decreased from 0.52 to 0 mg/ml. In contrast, as the cola concentration increased from 60% to 97%, casein concentration in the supernatant increased from 0 to 0.19 mg/ml. Fig. 3b and 3c show the supernatant turbidity of cola/casein mixtures containing different concentrations of cola. The supernatant turbidity was low in the presence of 30–60% cola at pH 3.2–3.6, and the ratio of the supernatant turbidity at pH 3.2–3.6 to that at pH 7.0–7.4 decreased to values below 0.1. As in the cola/milk system (Fig. 2), casein concentration in the supernatant first decreased and then increased as the cola concentration increased (Fig. 3a). These data indicated that RC occurred in the cola/casein system, in which milk is reduced to its major component, casein.

**Fig. 3.**
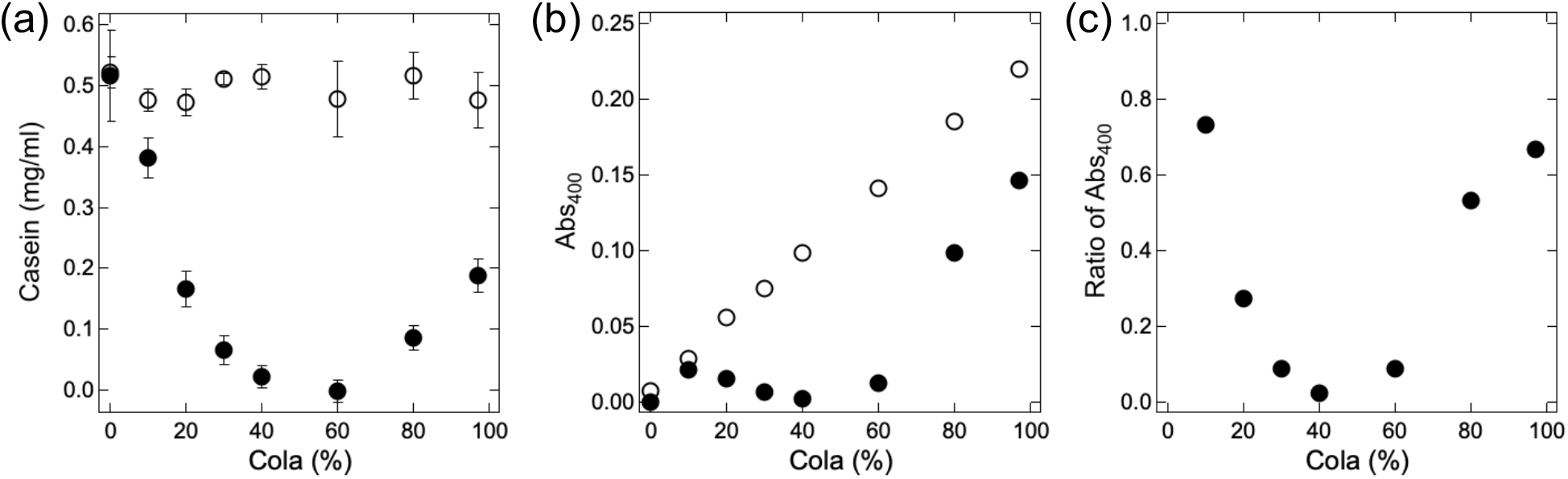
RC in the cola/casein mixture. (a) Casein concentration in the supernatant of mixture comprising 0.5 mg/ml casein and different concentrations of cola at pH 3.2–3.6 (closed circles) and pH 7.0–7.4 (open circles). Averages and standard deviations of triplicate measurements are shown. (b) Supernatant turbidity of mixtures containing 0.5 mg/ml casein and different concentrations of cola at pH 3.2–3.6 (closed circles) and pH 7.0–7.4 (open circles). (c) Ratio of the supernatant turbidity at pH 3.2–3.6 to that at pH 7.0–7.4 in the cola/casein mixtures.

Then, polyP was selected as a component of cola, where it is used as a food additive. To understand the phase behavior of a tetraP/casein mixture, we investigated the effects of the tetraP concentration on the casein concentration and zeta potential in the supernatant of tetraP/casein mixtures. Solutions were prepared using a constant concentration of casein (0.5 mg/ml) and varying concentrations of tetraP (0.01–2 mM) at pH 3.2–3.6. Fig. 4a shows casein concentration in the supernatant of tetraP/casein mixtures containing different concentrations of tetraP. Casein concentration was determined using the Lowry method. As tetraP concentration increased from 0.01 to 0.2 mM, casein concentration in the supernatant decreased from 0.39 to mg/ml. In contrast, as tetraP concentration increased from 0.2 to 1 mM, casein concentration in the supernatant increased from 0.01 to 0.40 mg/ml. Fig. 4b shows the supernatant zeta potential of tetraP/casein mixtures containing different concentrations of tetraP. As the tetraP concentration increased from 0.01 to 1 mM, the supernatant zeta potential decreased monotonically from 26 to − 23 mV, indicating charge inversion. As in the cola/milk (Fig. 2) and cola/casein (Fig. 3) mixtures, casein concentration in the supernatant decreased and then increased with the tetraP concentration (Fig. 4a). These data indicated the occurrence of an RC phenomenon associated with charge inversion of the tetraP/casein complex.

**Fig. 4.**
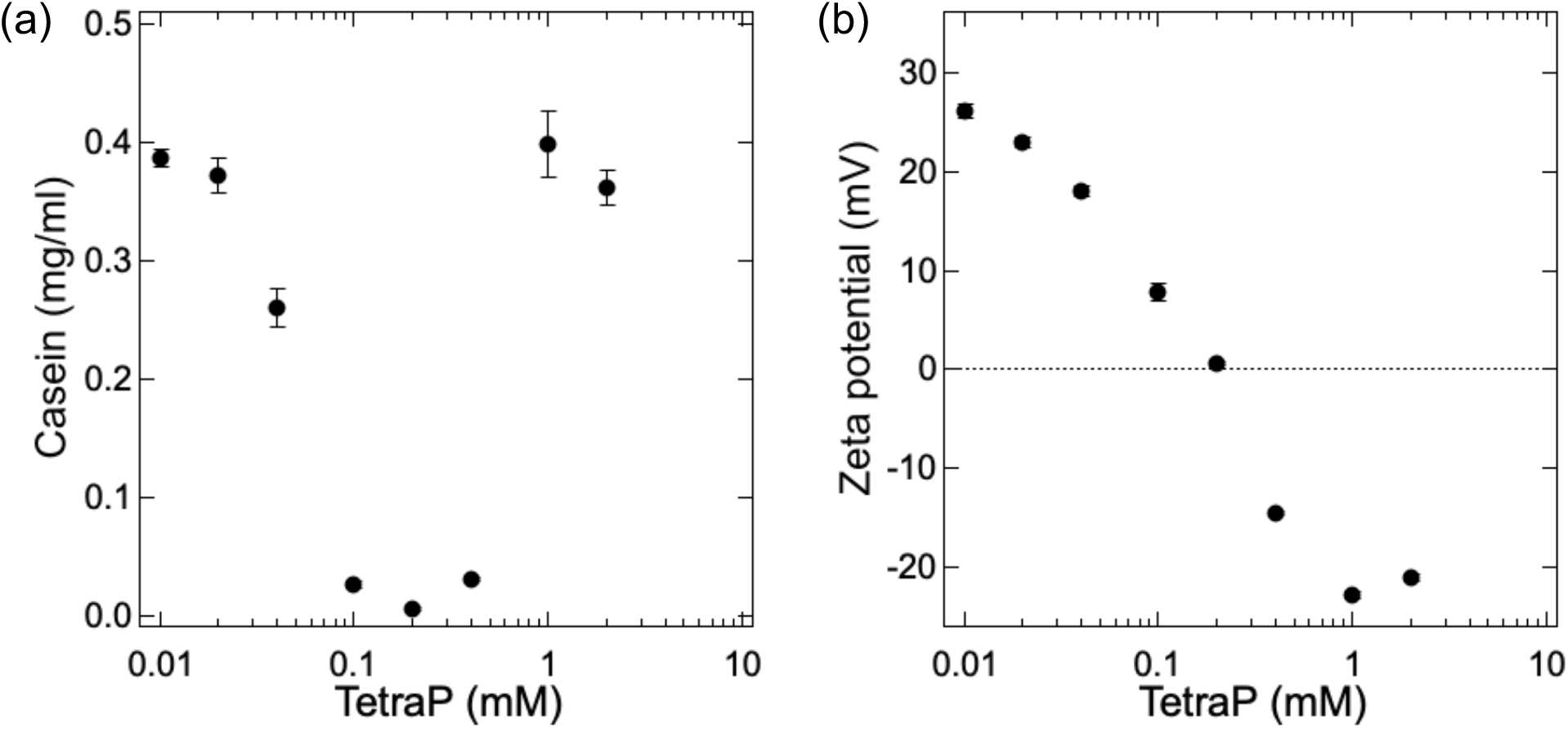
RC and charge inversion of the tetraP/casein mixture. Casein concentration (a) and zeta potential (b) in the supernatant of mixtures containing 0.5 mg/ml casein and 0.01–2 mM tetraP at pH 3.2–3.6. Averages and standard deviations of triplicate measurements are shown.

To clarify the effects of the polyP length on the phase behavior of polyP/casein mixtures, we investigated the impact of a concentration of tetraP, triP, or diP at different pH on the condensed or soluble state of polyP/casein mixtures. Solutions containing a constant concentration of casein (0.5 mg/ml) and varying concentrations of tetraP, triP, or diP (0.1–4 mM) were prepared, and the pH was adjusted by adding phosphoric acid. Fig. 5a shows the phase diagram of casein in the presence of tetraP at different pH. In the presence of 0.1 mM tetraP, casein condensed at pH 3.0–4.5, but not at pH values below or above this range. In the presence of 0.2–1.0 mM tetraP, casein was prone to condense at lower pH values. In contrast, casein did not condense at pH values above its p*I* (4.6) regardless of the tetraP concentration tested. Interestingly, the condensation of casein at its p*I* with 0.1 mM tetraP was suppressed in the presence of tetraP at a concentration of 0.2 mM or higher. These data indicated that tetraP shifts the pH range at which casein condensation occurs toward more acidic values. Fig. 5b shows the diagram of the condensed and soluble states of casein in the presence of triP at different pH. In the presence of 0.1 mM triP, casein condensed at pH values of 3.8–4.3, this range was narrower than that in the presence of tetraP. The pH range at which casein condensation occurred broadened toward lower values with increasing triP concentrations (above 0.1 mM). Casein did not condense at pH above its p*I* for all triP concentrations tested. We also constructed a phase diagram of casein condensation in the presence of diP (Fig. 5c). For all diP concentrations tested, casein condensed at pH 4.3– 5.0, which were values close to casein p*I*. Unexpectedly, diP, at least at the concentrations tested, did not affect the pH values at which casein condensation occurred, indicating that the negative charge of diP was not sufficient to modify the pH range for casein condensation.

**Fig. 5.**
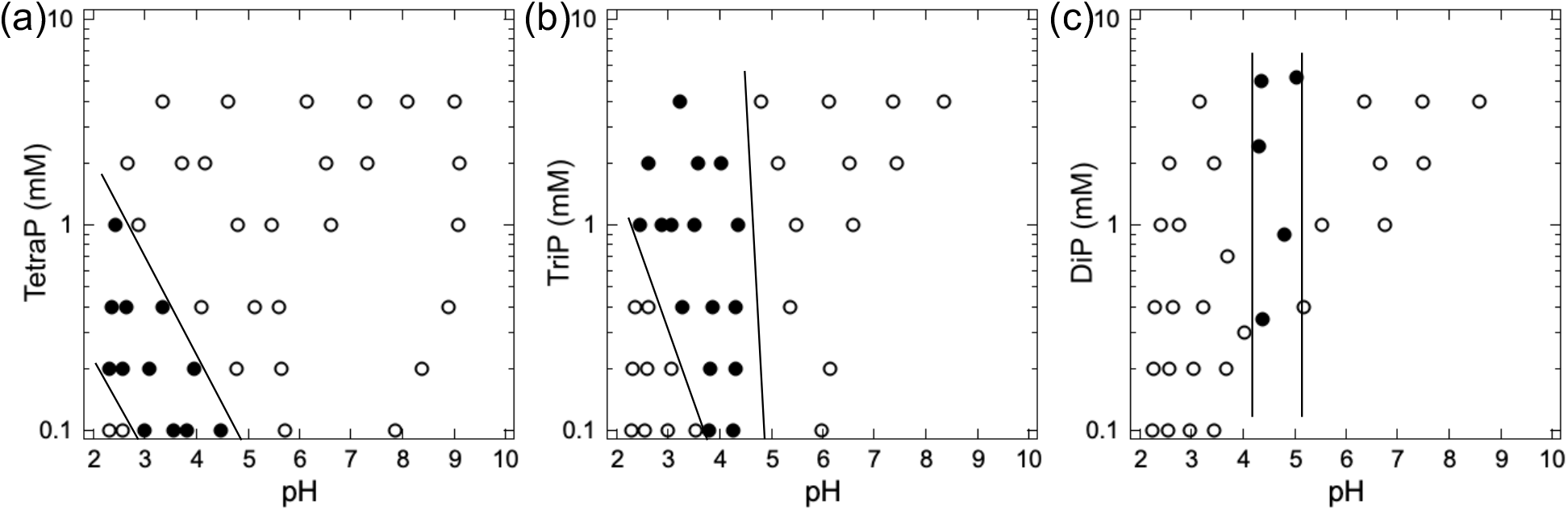
Phase diagram of casein condensation in the presence of (a) tetraP, (b) triP, and (c) diP. The solutions contained 0.5 mg/ml casein and 0.1–4 mM polyP at pH 2–10. Closed circles, condensed state; open circles, soluble state.

## 4. Discussion

Here, we generated a multicomponent complex system exhibiting RC behavior using cola and milk. At pH 3.2–3.6, condensation occurred only for cola/milk mixtures containing 30–40% cola but not for those with lower or higher cola content. This is a phase behavior typical of RC, where a system condenses only at moderate concentrations of multivalent ions or polyelectrolytes [33]. In contrast, at neutral pH (pH 7.0–7.4), there was no condensation at any tested concentration of cola. We then focused on casein, which constitutes 2.6% of milk and 79.5% of milk proteins. Casein has a positive charge at pH 3.2–3.6 and a negative charge at pH 7.0–7.4 because its p*I* is 4.6. There was RC in the cola/casein mixture as in the cola/milk system.

Quantification of the casein concentration in the supernatant revealed that casein was present in the condensate. Previous studies have shown RC of negatively charged proteins in the presence of multivalent cations [33–36]. Since casein has a positive charge at pH 3, multivalent anions in cola might have induced RC of casein. Therefore, we focused on polyP, a multivalent polyanion present in cola as a food additive [26]. The charge numbers of tetraP, triP, and diP at pH 3.4 determined by acid titration were − 4.0, − 2.9, and − 1.9, respectively (Fig. S1). As expected, RC occurred in the tetraP/casein mixture. Furthermore, the zeta potential changed from positive to negative as the concentration of tetraP increased, indicating charge inversion, a phenomenon typical of RC. Similarly, a system composed of the positively charged lysozyme and triP, which has a charge number of − 4.8 at pH 9, undergoes RC associated with charge inversion [30].

Since we found polyP important for the RC of casein, polyP is likely the main factor causing the RC of the cola/milk mixture. Cola contains approximately 0.20 mg/ml phosphorus [37], which corresponds to 6.5 mM phosphate units and is of the same order of magnitude as the concentrations used in the present study. In the production of cola, the addition of 300–1500 ppm polyP (4–19 mM phosphate units) is preferred for maintaining carbon dioxide bubbles [26]. However, because cola is a multicomponent system, components other than polyP may also contribute to RC. Cola contains small amounts of monovalent or divalent ions such as 0.02 mg/ml sodium, 0.02 mg/ml calcium, and 0.01 mg/ml magnesium [38]; however, several millimolar of these monovalent or divalent cations do not induce RC of BSA [33] and silica nanoparticles [8]. Caramel color is also a major component of cola and has a negative charge at pH 3.2–3.6 because its p*I* is below 2.5 [39]. It may condense with positively charged casein, resulting in the clear appearance of the supernatant. However, the RC behavior of the tetraP/casein mixture was very similar to that of the cola/casein and cola/milk systems. Therefore, polyP, with positively charged casein, is likely the major factor causing the RC in the cola/milk system. Thus, we have established a multicomponent complex system exhibiting RC using cola and milk. Moreover, RC in this system was fully reproduced by reducing the system to two components, i.e., polyP and casein.

The condensation of casein induced by polyP is explained by the following mechanism. The condensation of casein depended on the length and the concentration of anionic polyP, suggesting that it is induced by electrostatic interactions. At acidic pH below its p*I*, casein has positive charges and polyP has negative charges. Thus, positively charged casein is crosslinked by anionic multivalent polyP as it is observed in the PPC [5, 40] or the co-aggregation of two types of oppositely charged proteins [41, 42]. Furthermore, the chain length of polyP had an impact as casein condensed with 0.2 mM tetraP and 1 mM triP at pH 2.6, whereas it did not with diP. Additionally, 2 mM tetraP solubilized casein. These results indicated the importance of multivalent interactions between casein and polyP. When increasing polyP concentrations at pH below the p*I* of a protein, the system undergoes the transitions between the following states (Fig. 6): (i) a soluble one-phase state in which casein molecules have repulsive positive charges, (ii) a two-phase state with condensation in which positively charged casein is crosslinked by anionic multivalent polyP due to charge neutralization, and (iii) a soluble one-phase state in which an excessive amount of polyP increases the repulsion between casein molecules due to charge inversion in casein. The formation of soluble PPC has a similar mechanism, i.e., aggregative PPC is formed in the presence of low concentrations of polyelectrolytes and redissolves with high concentrations of polyelectrolytes [5, 40]. Therefore, our data suggest that multivalent ions or polyelectrolytes with high charge densities tend to induce RC.

**Fig. 6.**
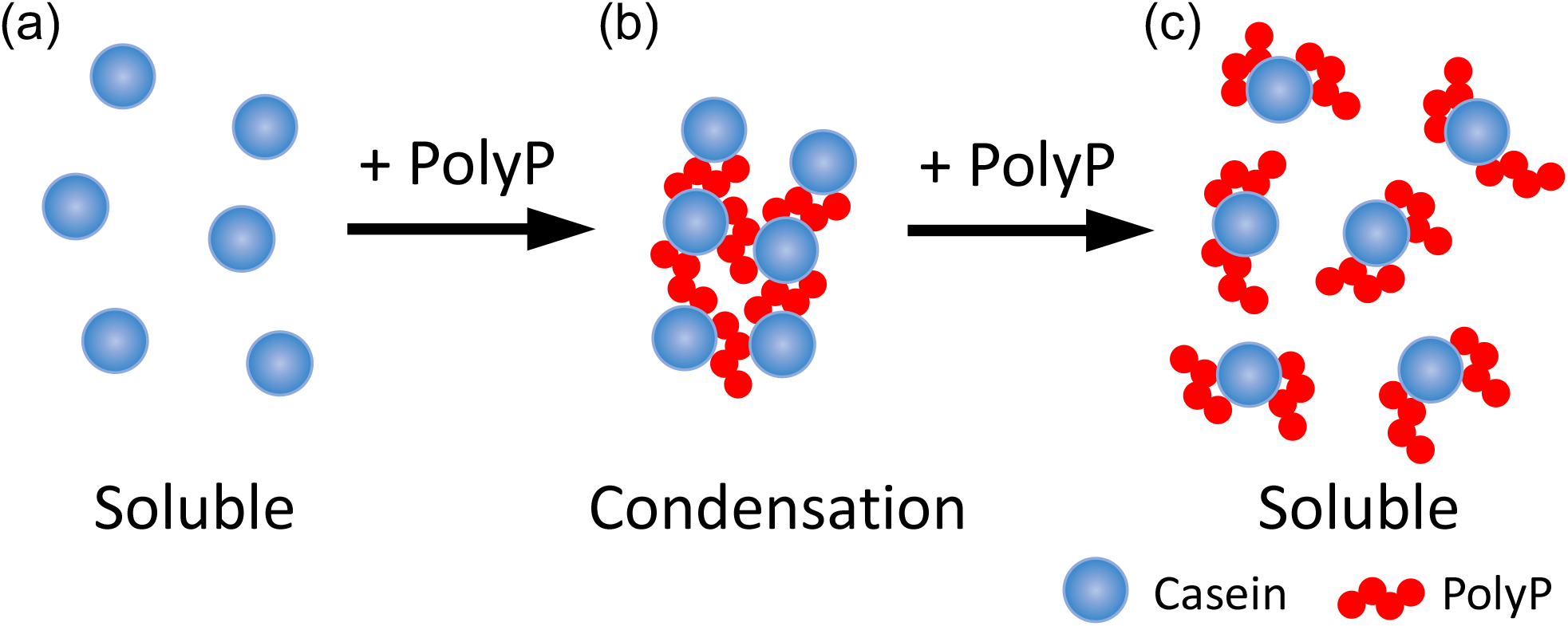
Schematic model of the RC of the polyP/casein mixture at pH below the p*I* of casein. (a) In the presence of low polyP concentrations, positively charged casein is soluble due to electrostatic repulsion. (b) With moderate polyP concentrations, positively charged casein is crosslinked by anionic multivalent polyP due to charge neutralization. (c) In the presence of high polyP concentrations, the excess of polyP increases the repulsion between casein molecules due to charge inversion of casein.

Condensed biomolecules, called “complex coacervate” or “condensates,” are known to play important roles in biological processes [2, 3, 43]. In living cells, such condensed states are called “membraneless organelles” (MLOs). MLO components can be exchanged with those in the surrounding compartments and are grouped [3]. Typically, they consist of charged biomolecules such as proteins, nucleic acids, ATP, and polyP [23, 44– 46]. Although the mechanism of MLO formation has been well studied, that of MLO dissolution is poorly understood. RC is one of the candidate mechanisms that explain both the formation and dissolution of MLOs. Some combinations of charged biomolecules, such as RNA/RNA-binding protein and RNA/positively charged peptide, exhibit RC behavior due to their high charge density [15, 18]. Hundreds of millimolars of monovalent or divalent salts can induce RC of intrinsically disordered proteins such as FUS, LAF1, and γD-crystallin [16, 17]. Recent studies have suggested a possible contribution of RC to biochemical timekeeping by generating condensates that form and dissolve as the concentration of transcription products monotonically increases [18]. However, RC has not been directly observed in biological cells because monotonically changing the concentration of only one component in a cell is almost impossible. Therefore, to understand the mechanisms of RC in complex systems such as the cytosol and food, it is important to construct a bottom-up model that includes multiple complex components. Here, we established a model multicomponent complex system exhibiting RC behavior. Moreover, RC was fully reproduced by reducing the system to two of its components. The use of such model systems will allow a deep understanding of the mechanisms of RC in multicomponent complex systems and thereby will contribute to elucidating the role of RC in biological systems.

## Acknowledgements

We are grateful to Dr. Yuji Goto (Osaka University) for valuable discussions. We are grateful for the kind support of Ishikawa Prefectural Nanao High School for the initial stage of this research and for the academic atmosphere of the University of Tsukuba, the University of Tokyo, and the Kumano Dormitory at Kyoto University. We thank Ms. Nanako Sakakibara (University of Tsukuba) for experimental support.

