## Supplementary material for "Reentrant condensation of a multicomponent complex system of biomolecules induced by polyphosphate": Figure S1

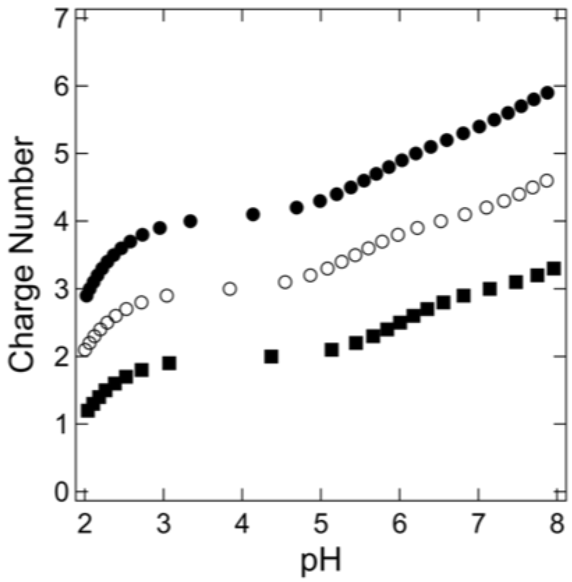


Figure S1. Impact of the pH on the charge number of polyP of different chain lengths determined by acid titration. Closed circles, tetraP; open circles, triP; and closed squares, diP.
